# Preserved geometry during representational drift enables stable perception and memory

**DOI:** 10.64898/2026.06.25.734656

**Authors:** Hajar Zaid, Evan S. Schaffer

**Affiliations:** Nash Family Department of Neuroscience and the Friedman Brain Institute, Icahn School of Medicine at Mount Sinai, New York, NY 10029, USA; CUNY Graduate Center, City University of New York, New York, NY 10016, USA

**Keywords:** representational drift, representational geometry, adaptive coding

## Abstract

In many brain regions, the stimulus tuning of neurons is stable on a timescale of hours but not on a timescale of weeks, a phenomenon often called ‘representational drift’. This would seem to imply that these brain regions cannot be used for stable recognition of sensory stimuli or the retrieval of associative memories learned several weeks prior. However, decoding approaches have demonstrated that in some cases, stable decoding of drifting representations is possible. In principle, adaptive decoding provides a plausible resolution to the paradox of how the brain operates with drifting representations, but we lack a deep understanding of what the requirements are for stable decoding to be possible. Here, we offer a general mathematical framework that explains when and why stable decoding from a drifting representation can be achieved. First, we demonstrate that both feedforward and recurrent networks preserve the geometry of their inputs when the network is sufficiently large, meaning that representational drift must also preserve geometry in these networks. Second, we demonstrate that drifting representations that have stable geometry are decodable with adaptive decoders. Therefore, not only the existence of preserved geometry in the presence of representational drift but also the ability to decode from drifting representations simply requires the population of neurons exhibiting representational drift to be large. This theoretical framework not only suggests that preserved geometry should be a general feature of drifting representations, it also explains the conditions under which empirical efforts to measure stable geometry will be successful.

## Introduction

In many brain areas, neural activity encoding sensory or task-related variables appears to gradually change over time, without a corresponding change in behavior. This phenomenon is commonly called “representational drift” (1–4). Representational drift on a timescale of 1-2 weeks has been observed extensively in the mouse brain in regions that are far from the sensory and motor peripheries, including CA1 and CA3 of the hippocampus (5–10), posterior parietal cortex (11) and retrosplenial cortex (12). Sensory cortices appear to be heterogeneous in their degree of stability, but representational drift has been observed in piriform cortex (13), primary auditory cortex (14), barrel cortex (15, 16), and visual cortex (17–20). How does the brain maintain stable percepts, memories and behaviors over long timescales if many brain regions exhibit drift on much shorter timescales?

One interpretation of the observation of representational drift is that brain regions exhibiting drift only maintain information on a timescale of weeks. This would imply that stable perception, the retrieval of longer-term memories, and the formation of longer-term behavior policies must be performed by other regions. This interpretation is problematic due to the sheer number of brain regions that exhibit representational drift. Instead, two different classes of computational models have been proposed to reconcile how stable behavior could be supported by drifting neural activity. The first possibility is that representational drift is in the null space of downstream regions, meaning that it has no effect on the information these downstream regions receive. This framework has proven to be valuable to explain other neural computations (21, 22), but the absence of evidence that representational drift is confined to specific dimensions of activity motivates the need for a second class of models in which downstream regions adapt to compensate for ongoing drift in their inputs.

A valuable insight into how different brain regions might adapt to ongoing drift is the possibility that the geometry of neural activity - the relationship between neural representations - is preserved, even when the representations themselves are changing. Observations across multiple brain regions suggest that geometry can in fact be stable despite change in representations due to a range of factors such as changes in internal state, context, and experimental variability (14, 17, 19, 23– 27). Network models have similarly demonstrated that representational drift driven by ongoing learning can preserve representational geometry (17, 24, 28, 29). However, it is not well understood whether there are general principles governing whether representational drift preserves geometry.

Adaptive decoding models that use prediction error-based learning can decode a one- or two-dimensional variable (e.g. an animal’s position in an arena) from drifting representations (24, 26). These observations demonstrate that prediction errors could provide a mechanism to compensate for drift. However, it is unclear whether this framework can be applied to higher-dimensional variables like object or odor identity: On any given day, an animal is unlikely to sample many of the dimensions of possible odors, meaning that the requisite prediction errors for adaptive decoding may be drastically under-sampled.

Here, we answer two open questions and demonstrate that the two are related: First, what are the sufficient conditions for representational drift to preserve geometry? Second, what are the sufficient conditions for stable adaptive decoding to be possible from drifting representations? We first show that changes in synaptic connectivity in either a feedforward or recurrent network produce representational drift but are guaranteed to preserve representational geometry, up to a bound that simply depends on the number of neurons in the drifting representation. Second, we demonstrate that preserved geometry is sufficient to enable an adaptive decoder to yield stable responses to unseen high-dimensional stimuli. Moreover, we demonstrate that a simple biologically plausible circuit can implement this adaptive decoding, maintaining an associative memory of unseen stimuli despite representational drift.

## Results

### Preserved input geometry with representational drift

We define the geometry of a representation as the set of all angles between pairs of stimulus representations (Fig. 1a). We begin by asking whether there are conditions under which representational drift is guaranteed to preserve geometry (Fig. 1b-c). We assume that representational drift is caused by synaptic changes, which could include both ongoing learning outside the control of an experimenter and probabilistic synaptic turnover. This assumption is consistent with the prevailing hypotheses for what causes representational drift (2, 4).

**Fig. 1.**
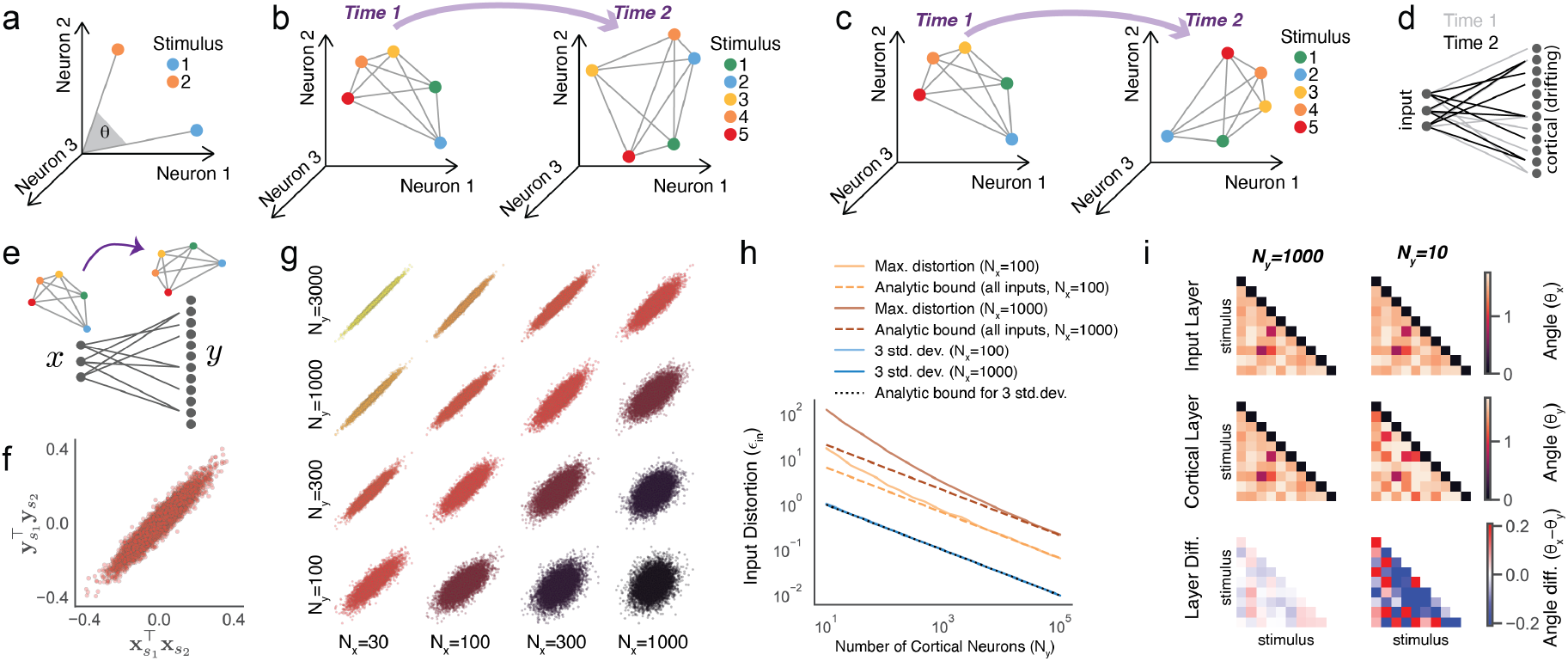
Feedforward neural networks preserve the geometry of their inputs. **a**. The set of all angles *θ* between pairs of stimulus representations defines the representational geometry. **b**. Schematic of drifting representations with drifting geometry. Each point represents the activity of all neurons to one stimulus. Note that from Time 1 (left) to Time 2 (right), the points are not only moving but also moving with respect to one another. **c**. Same as ‘b’ but demonstrating preserved geometry, whereby points do not move with respect to one another. **d**. A feedforward network in which activity in the input layer (**x**) is passed through synaptic connections that change over time to produce cortical activity (**y**). **e**. Schematic of preserved geometry from input **x** to output **y** for any fixed pattern of connectivity in a feedforward network. **f**. The similarity between random pairs of input patterns **x** and the corresponding similarity in the cortical patterns **y**. Each dot is the product of a unique pair of input patterns, with *N*_*x*_=100 and *N*_*y*_ =1000. **g**. Same as ‘f’ for a range of values of *N*_*x*_ and *N*_*y*_ . **h**. Two analytic bounds on distortion of input geometry (*ϵ* _*in*_), one that is accurate with high probability for random inputs (blue), and another that is exact and strictly true for all possible inputs (orange and brown). Each dashed line shows an analytic bound, and color-matched solid lines show results of simulations. **i**. Angles between pairs of stimulus representations in the input layer (top), cortical layer (middle), and the difference (bottom), for *N*_*x*_ = 100 and *N*_*y*_ = 1000 (left) or *N*_*y*_ = 10 (right).

In order to understand whether different patterns of connectivity between two brain regions produce the same representational geometry, it is instructive to first ask whether rep-resentational geometry is preserved from the first brain region to the second (Fig. 1e). If this were true for a random pattern of synaptic connectivity linking the two regions, it should be true for another random pattern of connectivity with the same statistics, meaning that changes in synaptic connectivity would not affect representational geometry. Thus, in order to show that geometry is preserved despite representational drift, it would be sufficient to show that geometry is preserved for any particular choice of connectivity. Therefore, we first ask whether random connectivity between two brain regions preserves representational geometry.

We first consider a simple two-layer linear feedforward network, but as we show below, our results generalize to models with biologically realistic nonlinearities and recurrent connectivity. The feedforward network (Fig. 1d) is given by

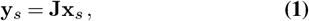

where 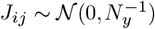 and **x**_*s*_ is the response to stimulus *s* in the input layer of the network, which for simplicity we assume is linearly related to the input. This network projects the *N*_*x*_-dimensional representation **x**_*s*_ to an *N*_*y*_-dimensional representation **y**_*s*_ (hereafter the “cortical layer”).

To examine representational geometry in each layer of the model, we compute the angles between the representations of any two stimuli, which are given by 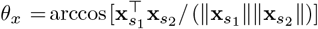 for the input layer (Fig. 1a). To examine whether geometry is preserved, we ask whether *θ*_*x*_ ≈ *θ*_*y*_ for all pairs of inputs, which we can simplify further by first examining whether 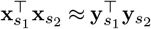. As shown in Figure 1f, the similarity of input patterns 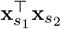 is highly correlated with the corresponding similarity of the output patterns 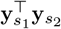 for *N*_*x*_ = 100 and *N*_*y*_ = 1000. We next asked how sensitive this correlation is to the number of neurons in the input layer *N*_*x*_ and the output layer *N*_*y*_. As shown in Figure 1g, the relationship between 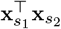 and 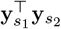 gets tighter as *N*_*x*_ decreases or *N*_*y*_ increases. In other words, input geometry is more accurately preserved in expansive networks where the output layer is much larger than the input layer, consistent with prior work (30–32).

To more rigorously understand how the preserved representational geometry depends on *N*_*x*_ and *N*_*y*_, we define *E*_*in*_ as the additive distortion from input geometry to cortical geometry,

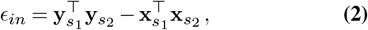

for any input patterns 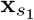 and 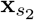 . We derive two bounds on *E*_*in*_, one that is accurate with high probability for randomly chosen input patterns and another that holds for all input patterns. To derive a bound on the distortion of random input patterns, we ask what the probability is of observing a given distortion *E*_*in*_. As detailed in Methods, a bound that captures *± d* standard deviations of the probability distribution *P* (*E*_*in*_) is

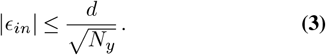

As shown in Fig. 1h, this bound is in good agreement with simulations, meaning that for large *N*_*y*_, representational geometry will be very well preserved.

The bound in Equation 3 is accurate with high probability, but it does not guarantee that geometry will be preserved for all possible input patterns. However, as detailed in Methods, a bound on *E*_*in*_ for all inputs is given by

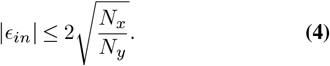

This bound applies to all possible inputs, meaning that if *N*_*y*_ is sufficiently large relative to *N*_*x*_, *E*_*in*_ is guaranteed to be small. This bound is loose for randomly sampled inputs, and it is only saturated when 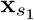 and 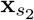 are both equal to the leading eigenvector of **J***T***J**. Therefore, we focus on the probabilistic bound given in Equation 3 in what follows. However, from both bounds, we can conclude that if and only if *N*_*y*_ is large relative to *N*_*x*_, the angles between stimulus representations are preserved from the input layer *x* to the cortical layer *y* (Fig. 1i, S1).

### Preserved input geometry in a recurrent network

The network we have considered thus far ignores recurrent connectivity within the cortex, but many of the brain regions that exhibit representational drift are highly recurrent. Therefore, we next ask whether the properties of preserved input geometry differ in a recurrently connected network (RNN, Fig. 2a). We consider a standard RNN of the form

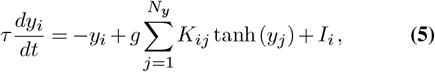

where 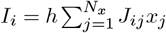 is an external input that is unique for each neuron, and each element of the matrices **K** and **J** is a Gaussian random variable with zero mean and variance 1*/N*_*y*_. Classic results demonstrate that a network of this form exhibits a phase transition from a fixed point of inactivity to chaotic activity in the large *N*_*y*_ limit when *g >* 1 in the absence of external input (33), but external input can suppress chaotic activity and drive the network to a stable fixed point (34), which we reproduce in Figure 2b for the case of static external input. For our purposes, the critical observation is that in both the *g <* 1 regime and the *g >* 1 regime with input-suppressed chaos, steady-state neural activity changes approximately linearly with respect to input magnitude *h* (34), which we reproduce in Figure 2c.

**Fig. 2.**
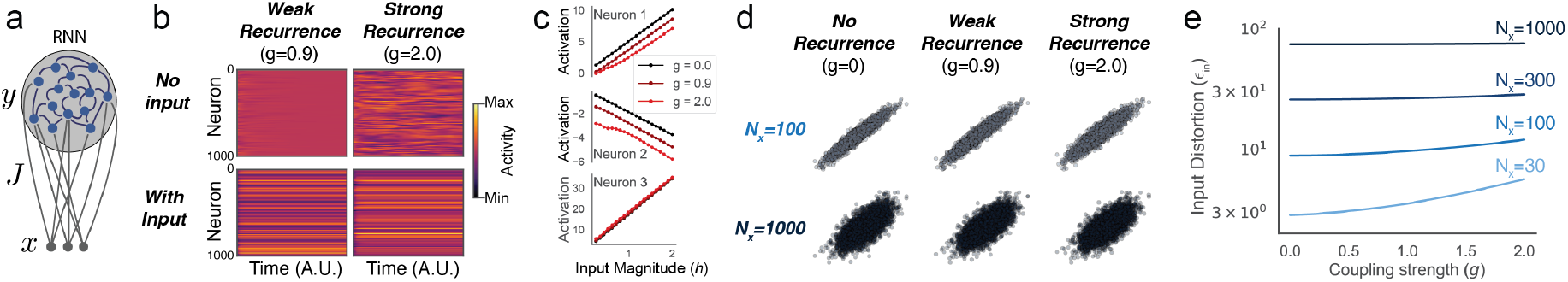
Recurrent neural networks preserve the geometry of their inputs. **a**. A recurrent network with external input. **b**. In the absence of external input, the network has a phase transition to chaos when *g >* 1 (strong recurrence), but both weakly and strongly recurrent networks exhibit stable responses to external input. **c**. Activity of 3 representative neurons versus input coupling strength (*h*) for different recurrent coupling strengths (*g*). **d**. The similarity between random pairs of input patterns **x** and the corresponding similarity in the steady-state cortical patterns **y**. Each dot is the product of a unique pair of input patterns, with *N*_*x*_=100, varying *N*_*y*_ and *g*. **e**. Distortion in input geometry (3 standard deviation bound on *ϵ* _*in*_, as in Fig. 1h) versus recurrent coupling strength (*g*) for *N*_*y*_ = 1000 and many values of *N*_*x*_.

The observation that input-driven activity in an RNN is approximately linear suggests that the input-driven activity of a recurrent network can be approximated by a linear feedfor-ward model, and therefore that the degree of preserved input geometry in an RNN should match that of a feedforward network. Consistent with this hypothesis, we find that input similarity 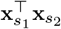 and output similarity 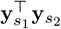 are largely in-dependent of coupling strength (*g*), spanning the range from a purely feedforward network (*g* = 0) to a strongly recurrent network, *g* = 2 (Fig. 2d-e). In general, this is not simply due to the external input dwarfing the recurrent input, as the recurrent input can be substantially larger than the external input (Fig. S1), and the input-driven response can differ substantially between the recurrent and feedforward case (Fig. 2c). These observations are therefore compatible with observations that recurrent inputs can dominate in cortex (35). Thus, for the remainder of this study, we examine the simple feedforward network, because it is sufficient for the questions of interest, without making assumptions about the impact of recurrent inputs in shaping activity in neural circuits.

### Stable geometry in drifting representations

How does preserved geometry from input to output translate into preserved geometry over time within the cortical layer? We model representational drift as random synaptic turnover in **J**. At every time step, a fraction *p* of all synapses are resampled from the distribution *J*_*ij*_ ∼ *N* (0, 1*/N*_*y*_). These changes to the connectivity matrix **J** cause the activity patterns **y**_*s*_ to drift (Fig. 3a). Intuitively, in the parameter regime for which input geometry is preserved at any time *t*, representational geometry in the cortical layer should be preserved despite representational drift. Validating this intuition, representational geometry is preserved despite representational drift for *N*_*x*_ = 100 when *N*_*y*_ = 1000 but not when *N*_*y*_ = 10 (Fig. 3b).

We define *E*_*t*_ as the distortion in representational geometry within the cortical layer from time 0 to time *t*,

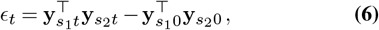

where 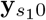 and 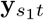 denote the cortical response to stimulus *s*_1_ at times 0 and *t*, respectively. Because *E*_*t*_ quantifies the distortion in geometry relative to time 0, and **J**_0_ ≈ **J**_*t*_ for small *t, E*_*t*_ is a growing function of time (Fig. 3c). However, the growth over time of *E*_*t*_ is bounded and saturates when the two matrices **J**_0_ and **J**_*t*_ are fully decorrelated (Fig. 3c). In this asymptotic limit, the error induced on the relationship between 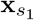 and 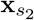 by **J**_0_ and **J**_*t*_ will be uncorrelated. Thus, in this decorrelated limit, the variances of the two single-time errors add, meaning 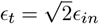. Therefore, a bound for the drift in the geometry of random inputs is given by

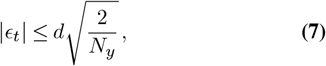

where *d* corresponds to the number of standard deviations to include in the bound on typical input patterns, as in Equation 3. As shown in Figure 3d, this analytic bound provides a good fit to the distortion in representational geometry observed in simulations. Therefore, geometry is preserved despite representational drift, as long as the brain region exhibiting drift has a sufficiently large number of neurons.

**Fig. 3.**
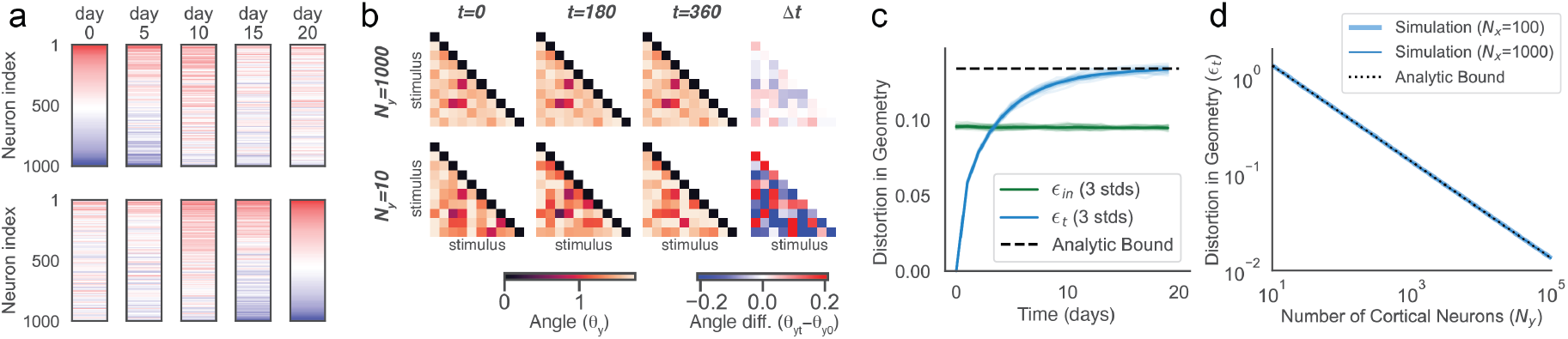
Representational drift preserves geometry. **a**. Drift in synaptic connectivity causes representational drift. Shown is **y**_*st*_, the response in the cortical layer of the network to a single input *s* as a function of time, sorted by response at *t* = 0 (*top*) or response at *t* = 20 (*bottom*) from max (red) to min (blue). **b**. Angles between pairs of stimulus representations in the cortical layer at *t* = 0, *t* = 180, and *t* = 360, for *N*_*x*_ = 100 and *N*_*y*_ = 1000 (*top*) or *N*_*y*_ = 10 (*bottom*). The difference between angles for *t* = 0 and *t* = 360 is shown at right. **c**. Comparing the representational geometry in the cortical layer at time 0 to time *t* defines the within-layer distortion (*ϵ* _*t*_) due to representational drift (blue). Distortion in representational geometry from the input layer to cortical layer (*ϵ* _*in*_) (green) and the analytic bound on asymptotic *ϵ* _*t*_ (dashed black) are shown for reference. **d**. Comparison between observed distortion in time (*ϵ* _*t*_) for two values of *N*_*x*_ and the analytic bound derived from *ϵ* _*in*_ (line from *N*_*x*_ = 100 hidden behind line from *N*_*x*_ = 1000).

### Stabilizing a memory by leveraging preserved geometry

Next, we ask how a biological circuit could take advantage of preserved geometry to maintain memories despite representational drift. Specifically, we ask whether preserved geometry offers a solution to the problem that an animal may be exposed to a limited set of stimuli on any given day and nevertheless needs to maintain stable responses to all possible stimuli. In principle, stable geometry implies that knowing how the representation of some stimuli have changed could be used to predict how others have changed. To test this intuition, we ask whether an adaptive readout can stabilize associative memories of arbitrary stimuli or merely those that are similar to ones that were recently experienced.

To study the impact of preserved representational geometry on learning, we introduce a linear readout whose response *z*_*s*_ to any stimulus *s* is a weighted sum of the response of neurons in the cortical layer, *z*_*s*_ = **w***T***y**_*s*_ (Fig. 4a). We consider a scenario in which the readout is trained on **y** * _0_, the cortical representation of one stimulus at time *t* = 0. Specifically, we implement a simple form of Hebbian learning (36) that sets the weights for the readout proportional to the activity produced by the stimulus being learned, **w**_0_ ∝ **y**_*0_. We then ask how well the readout can distinguish the trained stimulus from a panel of novel stimuli, and whether specificity for **y**_\**t*_ can be stabilized without having to experience the target pattern **y**_\**t*_. We quantify the selectivity of a readout as the signal-to-noise ratio for the learned stimulus (Methods).

**Fig. 4.**
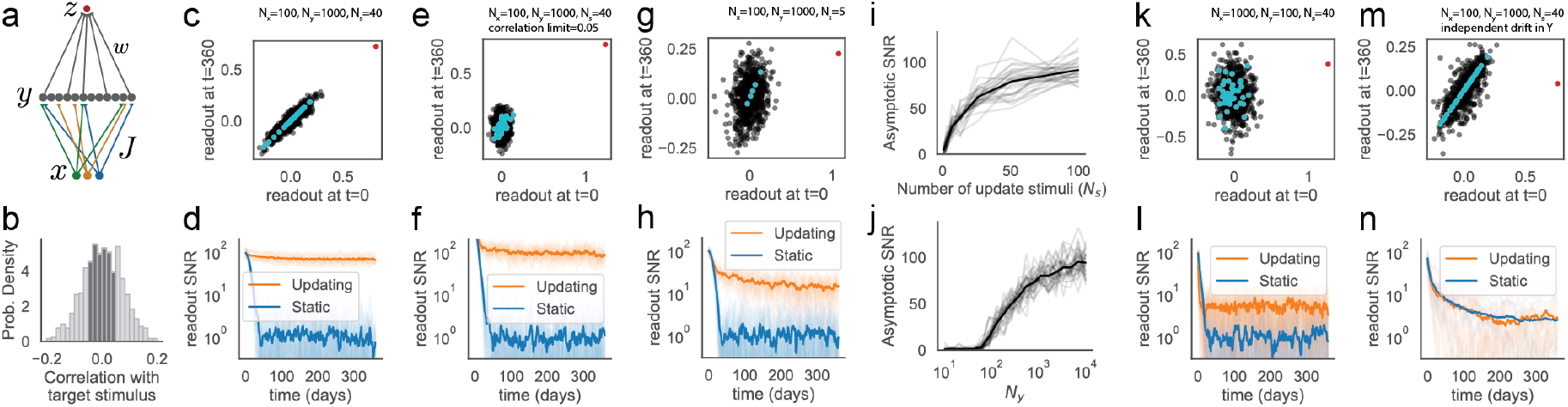
Preserved geometry in drifting representations enables stable memory. **a**. A feedforward network with trained linear readout *z*. **b**. After initial training, the readout weights **w** are updated using random stimuli **x**_*s*_ whose correlations with **x**_*_ are small. The full distribution of correlations is shown in light/dark gray, whereas only the stimuli producing correlations less than *±* 0.05 (dark gray) are used in ‘e-f’. **c**. The readout’s response to the target stimulus **x** (red), the most recently experienced stimuli **X** (cyan), and other stimuli (black) for time 0 versus time *t* = 360, for *N*_*x*_ = 100, *N*_*y*_ = 1000, and *N*_*s*_ = 40. **d**. Readout SNR over time for the same parameters as ‘c’, with (orange) and without (blue) updates to **w**_*t*_. **e-f**. Same as ‘c-d’, but restricting the stimuli **x**_*s*_ allowed for updating **w**_*t*_ to correlation magnitude no greater than *±* 0.05, as shown in ‘b’. **g-h**. Same as ‘c-d’, for *N*_*s*_ = 5. **i**. Readout SNR versus *N*_*s*_ for *N*_*x*_ = 100 and *N*_*y*_ = 1000. **j**. Readout SNR versus *N*_*y*_ for *N*_*x*_ = 100 and *N*_*s*_ = 40. **k-l**. Same as ‘c-d’, but for *N*_*x*_=2000. **m-n**. Same as ‘c-d’, but for independent drift in **Y**_*t*_ rather than **J**.

The readout is presented with a set of arbitrary stimuli, *S*, at every subsequent time step after initial training, where the number of stimuli in *S, N*_*s*_, is small. We allow feedback from the readout to update the weights **w**_*t*_ so as to adapt to stabilize the readout’s response to incoming stimuli, similar to other adaptive decoding models (24, 28). Specifically, we update **w**_*t*−1_ with a prediction error by comparing the desired output **z**_*S*,ref_ and the current output **z**_*St*_,

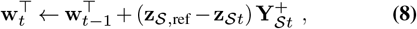

where 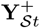 is the pseudoinverse of **Y**_*St*_, the matrix of responses of every neuron to every stimulus in *S*. We first take **z**_*S*,ref_ to be **z**_*S*0_, the readout’s response to any set of stimuli *S* at the time that the associative learning event **w**_0_ → **y**_*0_ occurred, and we begin by assuming these responses are known. We will relax this assumption below. This update rule can be seen as a form of Difference Target Propagation (37). Although this learning rule only explicitly stabilizes **z**_*St*_ for the observed stimulus set *S*, which does not include the target stimulus **x**_*_, we find that the SNR of the readout is very stable (Fig. 4c-d). Strikingly, this is despite the representation of every stimulus, including the target stimulus, changing entirely (Fig. 3a). In contrast, in the absence of the update rule for **w**_*t*_, the readout selectivity for **y**_*_ goes to zero because the cortical representations **y**_*st*_ are drifting (Fig. 4d).

The observed stimulus set *S* is selected randomly but never contains the trained pattern **y**_\**t*_. Nevertheless, one might imagine that maintaining the readout’s specificity for **y**_\**t*_ requires that some of the observed stimuli are similar to **y**_\**t*_. To test this, we asked whether specificity of the readout can be maintained if no observed stimulus during the maintenance period has a correlation magnitude greater than *±*0.05 with **x**_*_ (Fig. 4b). As shown in Figure 4e-f, the readout still achieves stable specificity while learning from this stimulus set, despite none of these stimuli being correlated with the target stimulus. However, requiring that every stimulus in the stimulus set **X**_*S*_ be exactly orthogonal to **x**_*_ prevents the read-out from stabilizing (Fig. S3). Thus, successful learning requires that the projection of **x**_*_ onto the sampled stimuli **x**_*S*_ is nonzero, but randomly selected **X**_*S*_ are sufficient despite their projection onto **x**_*_ being typically very close to zero.

Although maintaining specificity for **y**_*_ does not require observed stimuli to be correlated with **y**_*_, it depends on the number of observed stimuli. Readout performance is dramatically better when *N*_*s*_ = 40 than when *N*_*s*_ = 5 (Fig. 4g-h), but it degrades smoothly with *N*_*s*_, still achieving greater than chance performance when *N*_*s*_ is as small as 1 (Fig. 4i).

Intuitively, the ability to stabilize a trained readout using a prediction error from a small number of arbitrary stimuli should only be possible when drift preserves geometry. To test this intuition explicitly, we first simply increased *N*_*x*_ while keeping *N*_*y*_ constant, which distorts the input geometry (Fig. 1). As expected, this leads to poor readout performance (Fig. 4k-l). To see this another way, stable selectivity of the read-out requires *N*_*y*_ to be sufficiently large (Fig. 4j), which is the same condition for geometry to be preserved in the cortical layer. We next asked whether a readout can have stable SNR if the cortical representations **y**_*s*_ drift independently rather than as a consequence of drift in synaptic connectivity **J**_*t*_. Specifically, we add independent and accumulating Gaussian noise to each cortical neuron’s response to each stimulus *s* at each time step *t*. As expected, in these conditions the readout can stabilize the patterns it observes (**Y**_*St*_), but this does not generalize to the unseen target **y**_\**t*_ or other unseen inputs (Fig. 4m-n). Thus, maintenance of a stable readout of all possible stimuli using only a small number of samples requires preserved geometry.

### A direct/indirect circuit enables memory stability

In the previous section, we assumed that the plasticity rule for the readout has access to **z**_*S*0_. This memory of how the read-out would have responded to any possible stimulus *s* at time 0 is unrealistic. We next ask whether we can avoid requiring such a memory. Recall that the readout weights **w** are proportional to **y** * _0_ through Hebbian plasticity (36), meaning that the readout responses can be rewritten as 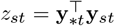, the projection of the representations of stimulus *s* onto the trained stimulus. From the observation that **x***T***x** ≈ **y***T***y** in Figure 1f, it follows that *z*_*st*_ can be approximated by 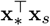. We incorporate this observation into our model by defining synapses **w**_*x*_ and **w**_*y*_ as the weights onto the readout from **x** and **y**, respectively (Fig. 5a), where both sets of synapses are subject to Hebbian plasticity during an associative learning event. This means that 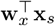 is equivalent to 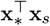. We again use the update rule for **w**_*y*_ given in Equation 8, but now with 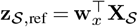.

**Fig. 5.**
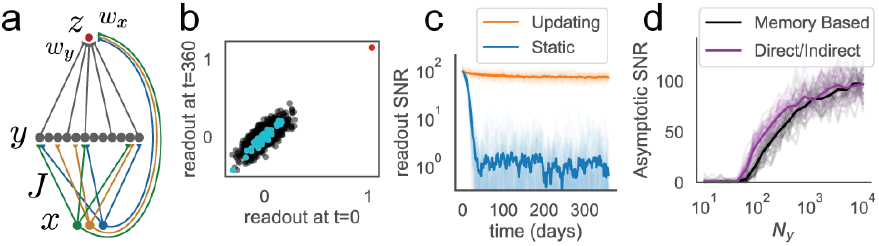
A direct/indirect circuit architecture enables memory stability. **a**. Model schematic. The readout receives input from both *x* and *y* via synapses *w*_*x*_ and *w*_*y*_ that obey the same Hebbian plasticity rule. **b**. The readout’s response to the target stimulus **x**_*_ (red), the most recently experienced stimuli **X**_*S*_ (cyan), and other stimuli (black) for time 0 versus time *t* = 360, for *N*_*x*_ = 100, *N*_*y*_ = 1000, and *N*_*s*_ = 40. **c**. Readout SNR over time for the same parameters as ‘b’, with (orange) and without (blue) updates to **w**_*t*_. **d**. Readout SNR versus *N*_*y*_ for *N*_*x*_ = 100 and *N*_*s*_ = 40 for this ‘direct/indirect’ architecture (purple). The comparable curve for the memory-based model (reproduced from Figure 4j) is also shown for reference (black).

We define the readout’s response to a given stimulus as 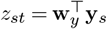, as for the models described above, meaning that the direct inputs from **x** only affect the plasticity of **w**_*y*_, not the output. This ensures that stable performance of the read-out is not simply driven by stable inputs that circumvent the drifting layer. As shown in Figure 5b-c, this readout that compares direct and indirect inputs achieves stable specificity for the trained input, comparable to and even slightly exceeding the performance of the original readout that was given access to **z**_***S***0_ (Fig. 5d). Therefore, a direct/indirect circuit architecture enables direct inputs from **x** to stabilize the readout’s responses to **y**. Biologically, this is also motivated by the pervasive direct/indirect architecture found throughout the brain on multiple spatial scales, including circuits exhibiting representational drift such as the piriform cortex (38) and hippocampus (39).

## Discussion

Stable behavior is often assumed to require stable neural representations. Here, we developed a theoretical framework to understand how stable behavior can arise despite representational drift, built on two linked ideas: bounds on the conditions under which drift preserves representational geometry, and an adaptive decoding scheme that can exploit that preserved geometry to maintain a stable readout. We have demonstrated that representational drift preserves geometry in expansive circuits, in which the downstream population is much larger than the upstream one. This result is consistent with earlier theoretical work showing that expansion supports generalizable representations (30–32, 40). This regime is also of particular interest because it is observed across many brain regions and species, including the mammalian and invertebrate olfactory and visual systems, the cerebellum, and the hippocampus, among others.

Our framework also suggests a simple way that down-stream circuits could leverage preserved geometry to compensate for drift. First, preserved geometry ensures that sampling a small set of arbitrary stimuli is sufficient to stabilize a readout of even a high-dimensional representation space. Second, the reference signal required for correction can be supplied through direct input from the preceding layer rather than requiring memory. Our theory is also broadly consistent with recent evidence that cortical learning may rely on target-propagation-like principles (41). Together, these observations suggest that a feedback controller could produce a stable output despite ongoing representational change in its input, only relying on mechanisms compatible with known circuit architecture. These results extend ideas from our previous work showing that different random projections of the same input ensemble can support similarly effective readouts, but only in sufficiently high-dimensional regimes (30).

Preserved geometry has been experimentally observed in primary motor cortex (23), primary visual cortex (17, 19, 27), medial entorhinal cortex (25), and hippocampal CA1 (26), but notably not in olfactory representations in piriform cortex (13). Our framework provides a simple explanation for why geometry would appear not to be preserved in piriform cortex: In contrast to the low-dimensional task variables examined in other brain areas, the dimensionality of odor identity may be upwards of 1000, the number of olfactory glomeruli in the mouse. Because the preservation of geometry depends on a ratio between this input dimensionality and the number of neurons in the drifting brain area, preserved geometry in piriform cortex would be barely discernible for arbitrary odors when 300 neurons are recorded, even if it were present. However, our theory predicts that preserved geometry in piriform cortex would become apparent with any of the following experimental modifications: Increasing similarity of the studied odors, decreasing the number of olfactory receptors, or increasing the number of recorded neurons (See Fig. S1).

A growing literature points to several candidate mechanisms for the cause of representational drift, including on-going learning, synaptic turnover, homeostatic plasticity, and synaptic volatility (1, 2, 4, 42–44). Computational models support the plausibility of each of these mechanisms, suggesting that drift can arise naturally from continual learning (45, 46), consolidation (47–50), and changes in neural excitability (51). Whether stable behavior ultimately requires stable activity in motor cortex remains somewhat controversial, but activity in motor cortex appears to be comparatively stable in the absence of behavioral changes (52, 53) (but see (54)). Taken together, these studies suggest that drift may be an unavoidable feature of adaptive systems, and consequently that adaptive decoding may be a critical function of many brain regions.

Our framework is deliberately agnostic about which of the potential causes of drift is most responsible in any given circuit. This is important because the causes of drift may vary across brain areas and timescales, whereas the geometric constraints on stable decoding may be considerably more general. However, one particularly compelling possibility is that drift depends on experience rather than simply on the passage of time. This is supported by observations of experience-dependence of place-cell drift in hippocampal CA1 (55, 56), orientation selectivity in visual cortex (20) and odor tuning in piriform cortex (13). The possibility that drift is driven by experience means that the number of stimuli needed to stabilize downstream decoding may always be satisfied, if drift slows down when fewer stimuli are experienced.

We have defined representational drift as a slow process driven by synaptic dynamics, but apparent representational change can also reflect behavioral variability, contextual modulation, arousal, or measurement noise. We anticipate that geometric principles similar to the ones we have developed here may also help reconcile how circuits maintain stable responses despite these changes in behavioral and internal states. The magnitude and structure of drift are also likely to differ across circuits and species. For example, in the primate inferior temporal cortex, many neurons maintain their selectivity for at least a month (57, 58), though some neurons are much more plastic (58). However, there is evidence for representational drift even in human primary visual cortex (27). Understanding how such differences relate to architecture, recurrence, redundancy, and behavioral demands will be an important direction for future work.

The broader implication of our results is that stable function does not require stable representations. What must be preserved is not the activity of individual neurons, but the geometry that allows downstream circuits to generalize and adapt to change. If that condition holds, then experience with a small number of inputs can be sufficient to infer and correct the displacement of an entire representational space. This framework is potentially compatible with standard engram-based theories of memory (59, 60), which hypothesize that certain dimensions of neural activity are stable. Moreover, extensions of the engram model to incorporate stochastic or fluid engrams have merged these ideas with the proposal that stable function emerges from dynamic engram membership (61).

Finally, identifying which neural computations can be stabilized by adaptive decoding is an important open question. The presence of representational drift in some cortical circuits may pose a constraint on the computations that can be maintained versus those that can only be performed transiently. Future work to identify these computational constraints has the potential to not only clarify our understanding of the consequences of representational drift, but also clarify our understanding of cortical computations more generally.

## Methods

### Notation

We denote matrices as capitalized and bold (**A**), vectors as lowercase and bold (**a**), and scalars as lowercase and normal font weight (a). We denote the elements of a matrix with indices and normal font weight (*A*_*ij*_). All vectors are assumed to be column vectors, so we denote row vectors as **a***T*.

### Feedforward Model

Let 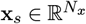 denote the input vector corresponding to the responses of *N*_*x*_ neurons to stimulus *s*, and let 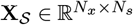 be the matrix whose columns are the responses **x**_*s*_ to each of the *N*_*s*_ stimuli in the stimulus set *S*, Each entry of **x**_*s*_ corresponds to the activation of one of the *N*_*x*_ neurons in response to stimulus *s*.

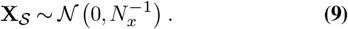

We model representational drift as random synaptic turnover in a connectivity matrix **J**_*t*_, where 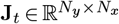 represents the connections from the input neurons to *N*_*y*_ neurons in the downstream layer of the network at the time step *t*. At each time step *t*, a fraction *p* of all synapses are resampled, and each new synapse is drawn independently from a normal distribution,

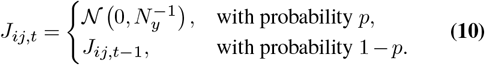

Consequently, the activity patterns **y**_*s*_ drift, as each **y**_*s*_ is a weighted sum of receptor activations with time-dependent synaptic weights,

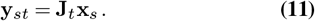

We define *z*_*s*_ as the output of a single downstream readout neuron receiving input from the cortical layer,

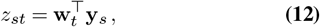

where **w**_*t*_ is a vector of connection strengths from each cortical neuron onto the readout *z*_*s*_.

### Update Rule for a Stable Readout

We define a target stimulus **x**_*_ and a corresponding cortical pattern **y**_*_. At time *t* = 0, we set **w**_0_ = **y**_*0_. The goal of maintaining selectivity to **y**_*_ would be achieved if 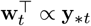, meaning that the weights would be maximally correlated with the target stimulus. However, this is inherently infeasible, as it would require that the synaptic update rule had continuous access to the original stimulus at all times *t*. Instead, we define a learning objective that minimizes the difference between the readout *z*_*s*_ at time 0 through time *t*, for a small number of stimuli *S*, which is achieved by updating **w**_*t*_,

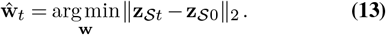

We let 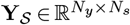 denote the response matrix corresponding to a subset of *N*_*s*_ stimuli, where each column represents the cortical response to one stimulus. The number of observed stimuli *N*_*s*_ is typically much smaller than the dimensionality *N*_*y*_ of the cortical representation. In this subset regime *N*_*s*_ *« N*_*y*_, the decoder must generalize from limited samples of the stimulus space. For the stimuli in *S*, the optimal **w**_*t*_ is achieved with the Moore-Penrose pseudoinverse, 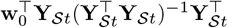. This pseudoinverse on its own does not result in a readout with stable performance due to the small number of stimuli in *S*. Instead, it serves as a correction term that we apply iteratively to adjust the weights at each time step *t*. We use an update rule inspired by Difference Target Propagation (DTP), where weight updates are driven by target-based corrections. The update follows:

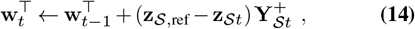

where 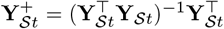. The target representation **z**_*S*,ref_ is set to either 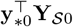, which is equivalent to **z**_*S*0_ or 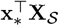. Under this update rule, the weights evolve as an accumulation of update vectors:

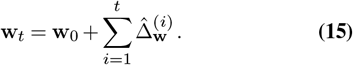

In order to evaluate how well DTP works to adjust the readout weights and generalize to all stimuli at time *t*, we define the Signal-to-Noise Ratio (SNR) of the readout’s selectivity for the target stimulus, *z*_\**t*_, as:

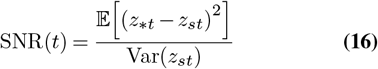

where the expectations are taken over all possible stimuli, not the set of stimuli used for updating, *S*.

### Recurrent Neural Network

To examine the degree of preserved input geometry in a recurrent network, we studied a model of the form

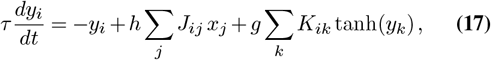

where the matrix elements *J*_*ij*_ and *K*_*ik*_ are chosen from a Gaussian distribution with zero mean and variance 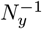. We exclusively studied the input-driven regime in which the dynamics converge to a fixed point, **y**_*s*∞_. To compare the geometry between inputs and steady-states in the network (Eq. 19), the recurrent network requires an additional normalization factor, 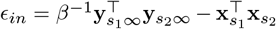. We estimate *β* numerically as 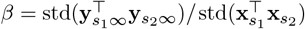, but a good approximation when *g* is small and/or *N*_*x*_ is large is *β h* (See Fig. S2).

Except where indicated otherwise, we take 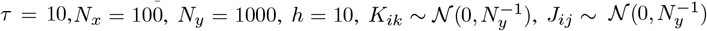, and each element of the input vector **x**_*s*_ is constant in time with a unique value for each neuron *i* and stimulus *s* chosen according to 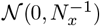, as in the feedforward network.

### Stable Geometry in Drifting Representations

To understand the conditions for drift to preserve geometry, we first find bounds for the distortion of input geometry in the cortical layer of the network. We then use this to find a bound on the distortion between two time points due to drift in the cortical layer. The condition for input geometry to be approximately preserved in the cortical layer is

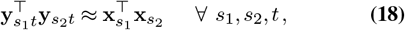

which we formalize by defining

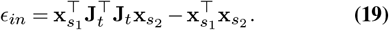

To define the worst case bound, it is useful to start with the condition for 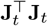 to be a frame for any individual input **x**_*s*1_, which generalizes the concept of a basis. A matrix **Φ** ∈ ℝ*n* is called a frame for R*n* if it satisfies the inequality:

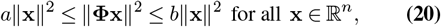

where 0 *< a < b* ≤ ∞. This condition ensures that **Φ** can only stretch or compress a vector a finite amount. For an input pattern **x**,

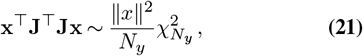

where 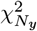 is the chi-squared distribution for *N*_*y*_ degrees of freedom. Plugging the expression above into the frame inequality,

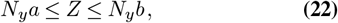

where 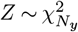. When *N*_*y*_ and *N*_*x*_ are large, but *N*_*y*_ *» N*_*x*_, the eigenvalue bounds of **J***T***J** (i.e., *a* = *λ*_min_, *b* = *λ*_max_) follow the Marchenko-Pastur Distribution:

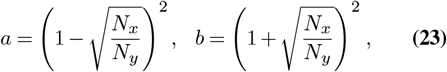

Although the Marchenko–Pastur bound is naturally written as a bound on the multiplicative distortion of the squared norm of a single vector, it also bounds the worst-case additive distortion of the inner product between two vectors. To see this, let *λ*_*i*_ be the eigenvalues of **J***T***J**. The largest possible squared-norm distortion is max_*i*_ | *λ*_*i*_ − 1 | . The largest possible additive inner-product distortion between two unit vectors is also max_*i*_ | *λ*_*i*_ − 1 |, because it is achieved by choosing both vectors to be the eigenvector corresponding to the most distorted eigenvalue. Thus, no pair of unit-norm inputs can have additive inner-product distortion larger than the worst single-vector squared-norm distortion. Therefore, defining *a* = 1 − *E*_*in*_ and *b* = 1 + *E*_*in*_, this implies that to first order,

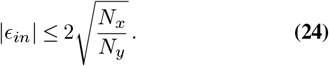

Thus, not only does **J** satisfy the frame condition, but in the large *N*_*y*_ limit, *E*_*in*_ approaches zero.

The bound above is true for all possible inputs, but it is not a tight bound for randomly drawn inputs **x**_*s*_. Instead of seeking a bound on all possible **x**_*s*_, we can derive the probability that the distortion exceeds a given *E*_*in*_. In high dimensions, most vectors are “typical,” and the probability of a random vector being atypical becomes vanishingly small.

The mean of *E*_*in*_ is zero. To compute the variance, let row *a* of **J** be 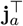, and define 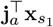 and 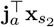 as *u*_*a*_ and *v*_*a*_, respectively. Then,

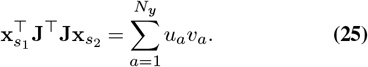

Conditioned on **x**_*s*1_ and **x**_*s*2_, the variables *u*_*a*_ and *v*_*a*_ are zero-mean jointly Gaussian with

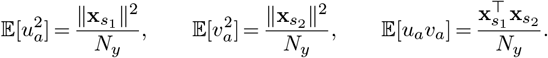

This means that

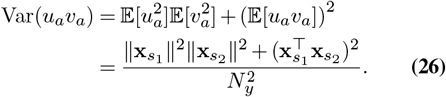

Because rows of **J** are independent, variances add across rows. Because 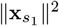 and 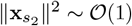 while 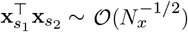, Var(*ϵ*_*in*_) ≈ 1*/N*_*y*_. If we define a typical inner-product error as one lying within *d* standard deviations of the mean, then:

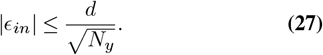

Finally, given the bounds for the distortion of input geometry described above, we can easily find a bound on the distortion between two time points due to drift in the cortical layer. A bound on the expected magnitude of the distortion from 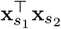 to 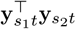 for typical inputs **x** is given by Eq. (27) for all times *t*. For two sufficiently different connectivity matrices **J**_0_ and **J**_*t*_, the corresponding single-time errors are approximately independent, so their variances add. Therefore, a bound on the distortion vector relating 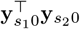 and 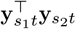 is simply 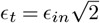.

## ACKNOWLEDGEMENTS

We are grateful to Larry Abbott, Carl Schoonover, Andrew Fink, Denise Cai, Peter Rudebeck, Frederic Stoll, Paul Kenny, and Richard Axel for helpful advice and comments, as well as the Simons Foundation (Simons Collaboration on the Global Brain NC-GB-CULM-00003215-07) for financial support.

## Supplementary Figures

**Fig. S1.**
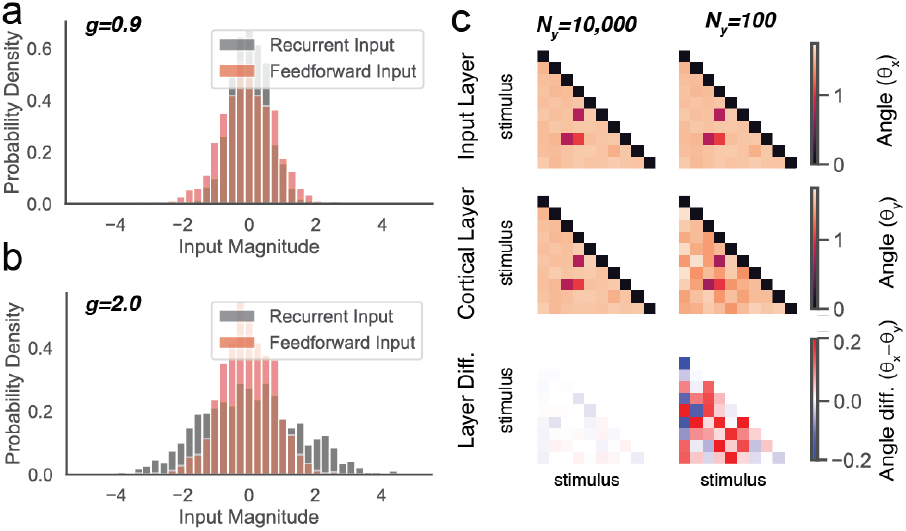
**a**. Distribution of input magnitudes onto recurrently connected neurons for recurrent and feedforward input, when *g* = 0.9. **b**. Same as ‘a’, for *g* = 2.0. **c**. Representational geometry in the input and cortical layers for 10 representative stimuli, as in Figure 1i, but with *N*_*x*_ = 1000 and *N*_*y*_ = 10000 or *N*_*y*_ = 100. Note that for *N*_*y*_ = 100, highly similar stimulus pairs (small *θ*) have preserved similarity in the cortical layer, whereas small differences between dissimilar pairs are lost in the noise in this regime.

**Fig. S2.**
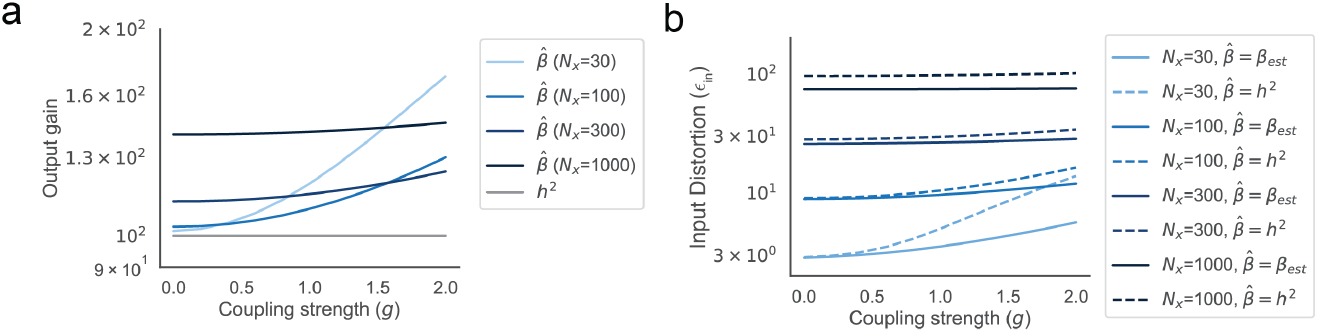
**a**. Comparing empirically estimated values of 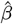 to *h*^2^ for a range of values of *g* and *N*_*x*_. **b**. Distortion in input geometry (3 standard deviation bound on *ϵ*_*in*_, as in Fig. 2e) versus recurrent coupling strength (*g*) for *N*_*y*_ = 1000 and many values of *N*_*x*_, comparing the results with both empirically estimated 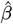 and setting 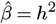.

**Fig. S3.**
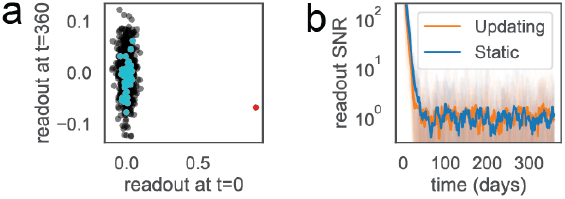
Same as Figure 4e-f, but for correlations between the sampled stimuli **X**_*S*_ and the target stimulus **x**_*_ forced to be zero. **a**. The readout’s response to the target stimulus **x**_*_ (red), the most recently experienced stimuli **X**_*S*_ (cyan), and other stimuli (black) for time 0 versus time *t* = 360, for *N*_*x*_ = 100, *N*_*y*_ = 1000, and *N*_*s*_ = 40. **b**. Readout SNR over time with (orange) and without (blue) updates to **w***t*.

## Notes

### Competing Interest Statement

The authors have declared no competing interest.

